# Dietary specializations are captured by jaw muscle proportions in mammals

**DOI:** 10.64898/2026.05.19.725803

**Authors:** Robert Brocklehurst, David M. Grossnickle, Joseph Bechara, Warren Cohen, Sharlene E. Santana, Christopher J. Vinyard, Andrea B. Taylor, Nicolai Konow

## Abstract

Mammalian diet and feeding ecology are often reflected by craniofacial skeleton specializations, but feeding requires skeletal actuation by a complex suite of muscles with varying sizes, lines of action, and mechanical function. While muscles play a critical role in feeding mechanics, and hence diet, it remains unclear how well variation in jaw muscle morphology predicts diet in mammals. We quantified the evolutionary interplay between mammalian muscle morphology and diet using a large and taxonomically broad sample. We measured the relative proportions and putative force production capacity, quantified as muscle physiological cross-sectional area (PCSA), for the major adductor complexes, along with a key jaw depressor, in 91 mammalian species (30 chiropterans, 33 primates, and 28 ungulates, carnivorans, rodents, and marsupials). We recovered clear dietary signals for several muscle complexes, with the medial pterygoid (larger in herbivores) and temporalis (larger in carnivores) performing best as dietary predictors. The medial pterygoid is particularly relevant for the mechanical innovation in mammals of moving the mandible along non-orthal, medio-lateral trajectories during mastication. Our findings underscore the intuitive, yet previously unquantified, importance of muscles in the evolution of mandibular roll, yaw, and lateral translation, all mammalian hallmarks of processing diverse types of food.

## INTRODUCTION

The ecomorphological paradigm states that organismal ecology is reflected by its morphology, which influences the ability of an organism to perform specific behaviors and, in turn, exploit ecological resources^1,2^. One example is the long-hypothesized link between dietary ecology and craniofacial morphology in mammals^3,4^. Feeding is a fundamental process of energy acquisition, and so selective pressures are expected to make feeding system morphology reflect the functional demands associated with feeding ecology. While general differences between skull and jaw shapes of herbivores and carnivores have long been established^3,5,6^, these ideas have generally evaded rigorous quantitative testing. More recently, morphometric advances have led to an upsurge in analyses addressing ecological traits — including diet — as macroevolutionary drivers of mammalian morphological change^7–9^. However, diet is far from always recovered as a strong predictor of skeletal morphology^10–12^. This is in part due to the skeleton being but one structural determinant of feeding system function. Other components of the feeding system - such as the muscles that drive skeletal motion and produce forces associated with feeding behavior - may better reflect ecological adaptations^13,14^.

In mammals, feeding involves extensive processing of food, where the mandible is moved by a suite of muscles relative to the skull around the temporomandibular jaw joint^15,16^. It is well established that mammalian jaw movements are three-dimensionally complex and, in addition to pitch rotations (orthal opening and closing), also involve yaw (mediolateral rotations) and translations in the fore-aft (propalinal) and transverse (side-to-side) planes^3,17–20^. Moreover, in many mammals, the lower jaw consists of two separate hemimandibles that can move relative to each other at their symphyseal midline^21–24^, and one or both mandible sides may rotate along their long axis^18,19,16^. These motions merge into an impressive diversity of ensemble movements of the lower jaw, ranging from relatively simple orthal cutting in mammalian carnivores^16,25,26^ to complex masticatory jaw motions for precise molar occlusion and food grinding in herbivorous mammals^18,27–29^. In turn, these movements are driven and controlled by the coordinated actions of a complex suite of jaw adductor (closer) and depressor (opener) muscles.

Differences in jaw motions involved in carnivory as compared to herbivory have been hypothesized to be associated with functional differentiation of the jaw adductor musculature; orthal cutting in carnivores is driven by the vertically oriented temporalis muscles, whereas transverse and proal grinding motions of mastication in herbivores are driven by the pterygoid and masseteric complexes^3^. The different functional roles of these muscles in the feeding system - their relative importance for different food processing behaviors, and thus, diets - are hypothesized to be reflected in the proportional size of each muscle or complex. However, this assumption remains untested, especially in a broad phylogenetic context. Whereas there is considerable variation in jaw muscle morphology across mammals, studies linking this soft-tissue diversity to differences in feeding behavior and/or diet have either been qualitative^3,4^, focused on specific mammalian orders ^30–35^, or on specific dietary groups ^36,37^. Consequently, we lack a broad comparative framework to determine how muscle morphology might reflect and predict diet across mammals as a whole.

Our aim is to measure the cross-sectional areas (CSAs) of the major jaw muscle complexes - temporalis, pterygoid, superficial masseter, zygomaticomandibularis/deep masseter, and digastric - in a broad phylogenetic suite of mammals. We take a functional approach to predicting muscle prominence and test the hypothesis that relative proportion (percentage of total cross-sectional area) of each jaw muscle complex reflects the evolution of dietary specializations. As carnivory (defined broadly here as including consumption of any animals, including insects) requires orthal jaw movements for cutting and crushing, we predict carnivores - the proposed ancestral mammalian diet^38^ - to have proportionally larger temporalis and deep masseteric complexes than omnivores and herbivores. Herbivory requires jaw movements in the transverse and/or proal directions for grinding when the molars are near occlusion. Hence, we predict herbivores to have proportionally larger superficial masseters and medial pterygoids. Muscle proportional sizes may also provide a functional perspective to characterizing omnivory, a dietary guild that historically has evaded characterization both ecologically^39^, and based on hard tissue morphology (e.g., compare omnivorous ursids to other carnivorans^40^). Evolution from a carnivorous ancestor into omnivory would require increased masticatory grinding of dietary plant matter, driven by the medial pterygoid and superficial masseter. We predict a gradual increase in the proportional size of these muscles going from carnivores, via omnivores to herbivores.

## RESULTS

### Muscle differences among diet categories

Data on craniofacial muscle cross-sectional area were gathered for 91 mammal species with a diversity of diets (Figure 1A) (Data S1). We calculated PCSA using muscle mass, fiber length and pennation angle. For sensitivity analyses on the importance of pennation, and how PCSA scales with muscle mass, see Supplemental Information. Dietary categorizations were based on the percentage of plant matter consumed by a species^6^ with supplemental evidence from other sources^41–43^ (Data S1). Associations were first tested by discretizing dietary percentage of plant matter into a dietary scheme with four categories (carnivory, 0–15%; carnivorous omnivory, 15–50%; herbivorous omnivory, 50–85%; and herbivory, 85–100%). To test for differences in muscle proportion between categories, we used phylogenetic generalized least-squares (PGLS) analyses, followed by analysis of variance (ANOVA), and post-hoc tests.

**Figure 1.**
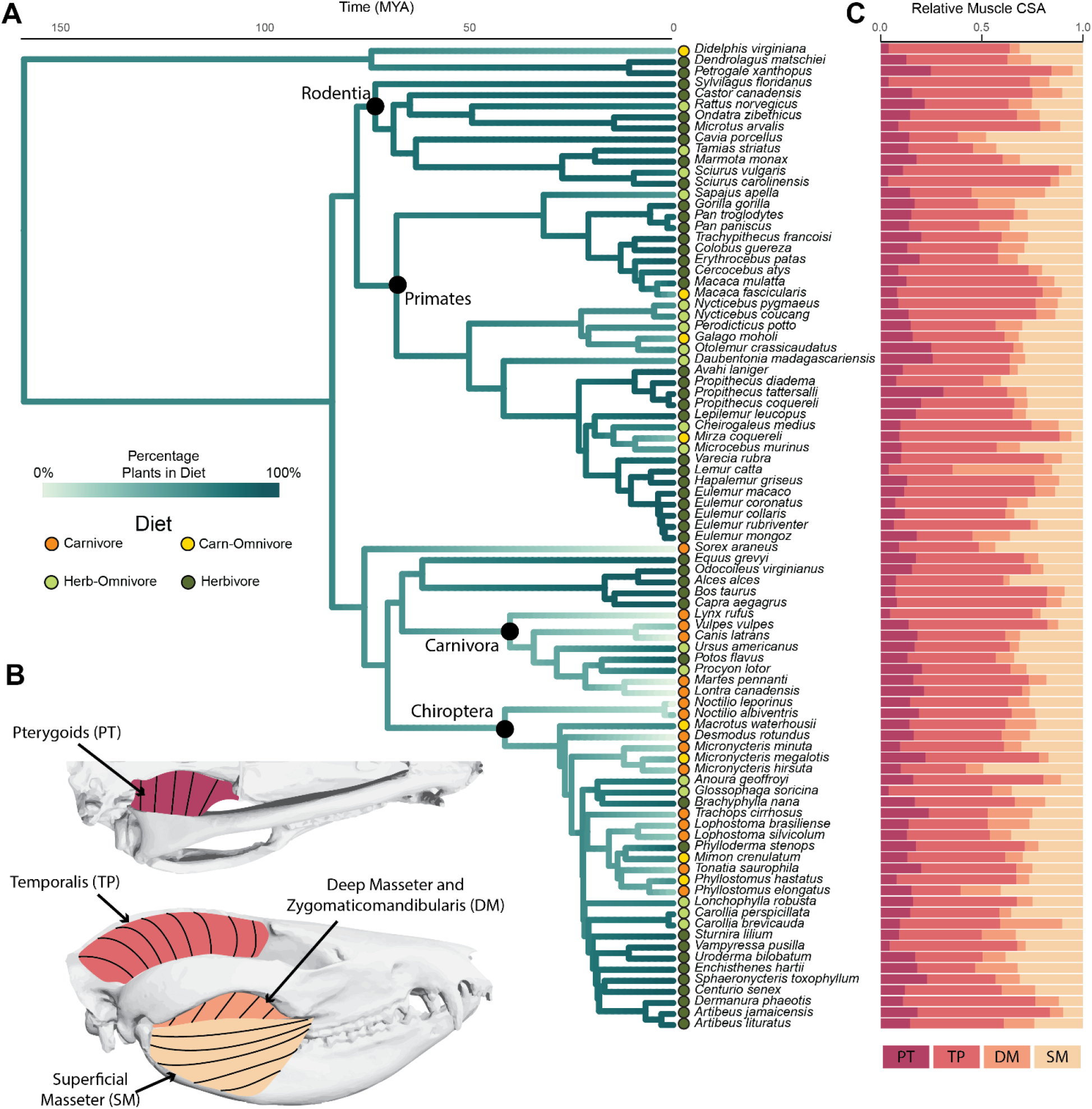
(A) Time-scaled phylogeny with dietary ancestral reconstructions and major clades labeled. Branch color coding reflects percentage dietary plant matter, with internal nodes being estimated based on continuous character mapping. Colored circles at the tips indicate diets of species in our dataset, coded as four discrete categories: carnivore (magenta, 0-15% plants in diet), carnivorous-omnivore (orange, 16-50% plant in diet), herbivorous-omnivore (51-85% plant material in diet) and herbivore (86-100% plant material in diet). (B) Jaw closer muscles sampled in this study: superficial masseter (SM), temporalis (TP), deep masseter and zygomaticomandibularis complex (DM) and pterygoid complex (PT) – in lateral (top) and ventral (bottom) views in *Didelphis virginiana* (Virginia opossum) (modified from Turnbull, 1970). (C) Stacked bar plots showing relative proportion of jaw adductor muscles (by proxy of cross-sectional area) in species from our sample.

Among the masticatory muscles, the pterygoids had the strongest relationship between muscle cross-sectional area and diet, being smallest in carnivores, followed by carnivorous omnivores and herbivorous omnivores, and largest in herbivores (Figure 2). Phylogenetic ANOVA recovered a significant relationship between pterygoid size and diet, and pairwise comparisons between diet categories indicated statistically significant differences between carnivores and herbivores (p < 0.001) and between carnivores and herbivorous omnivores (p < 0.05). Median PCSA for the temporalis was greatest in carnivores and consistently reduced in taxa classified as successively more herbivorous (Figure 2). Phylogenetic ANOVAs recovered a near-significant effect of diet on temporalis muscle PCSA, and a near-significant pairwise difference between carnivores and herbivores (Table S1). However, the relationship between the temporalis and diet was significant when using simpler diet classification schemes (two or three diets, see Supplemental Figures S2 and S3), and when using regular, non-phylogenetic ANOVAs (Table S2).

**Figure 2.**
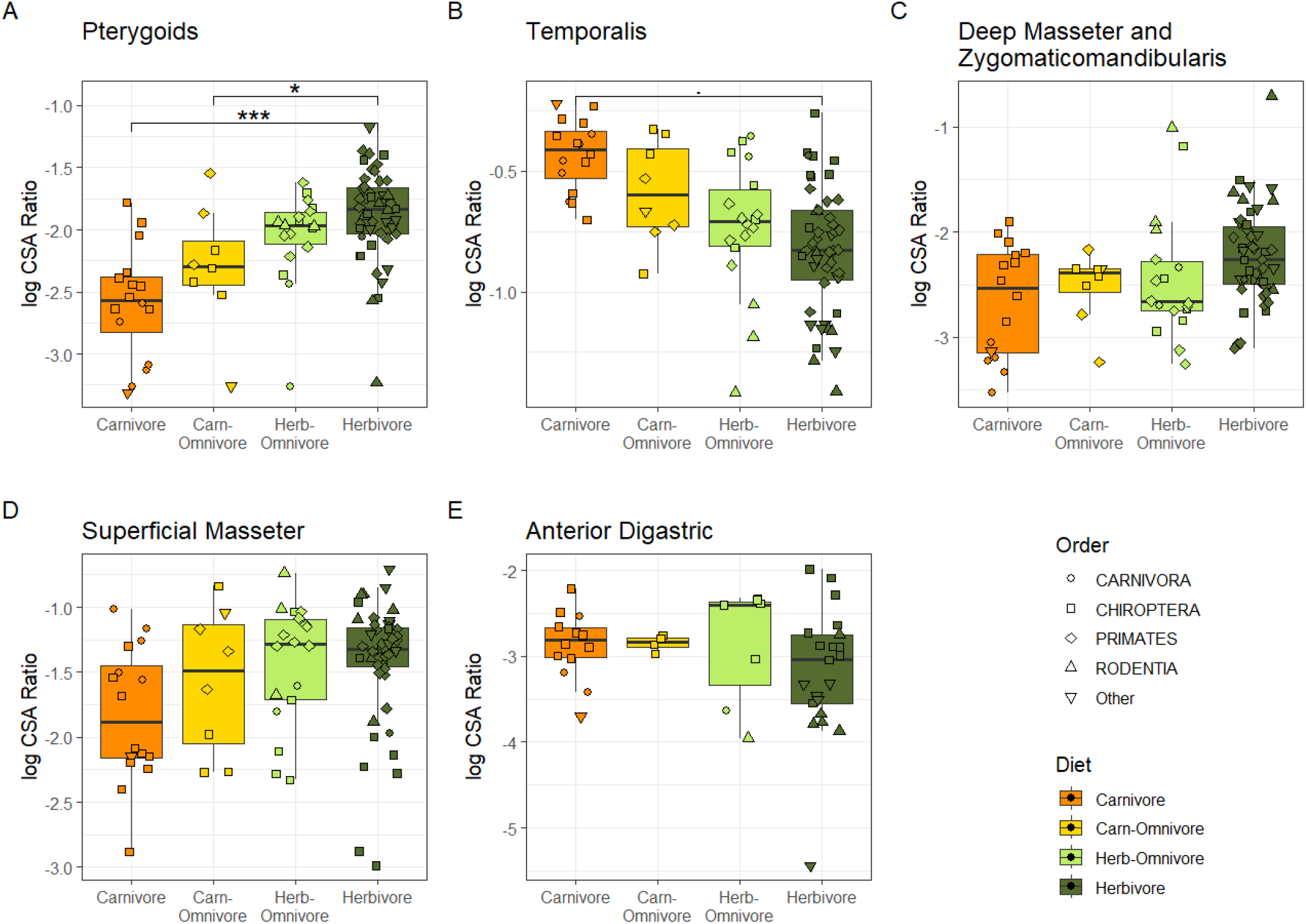
Differences in muscle proportions (separated in plot-space) by diet category. Diets (carnivory, carnivorous omnivory, herbivorous omnivory, and herbivory) are defined as in Figure 1. Muscles are the anterior digastric (AD), superficial masseter (SM), temporalis (TP), deep masseteric complex consisting of zygomaticomandibularis and deep masseter (DM) and pterygoid complex consisting of medial and lateral pterygoid (PT). Significant differences between diet groups, analyzed in a PGLS framework, are shown for the pterygoids and a marginally significant relationship is noted for the temporalis.

For the deep masseter and zygomaticomandibularis complex, herbivores showed relatively elevated median sizes compared to other groups, though with notable overlap, and some species were clear outliers. Phylogenetic ANOVA recovered no significant relationship between diet and deep masseter complex size, regardless of the dietary classification scheme used. Carnivores and carnivorous omnivores appeared to have lower median superficial masseter PCSA than herbivorous omnivores and herbivores, again with substantial data overlap (Figure 2B). Whereas superficial masseter size in herbivores versus carnivores was significantly different using non-phylogenetic tests (Table S2) there were no significant differences between groups when effects of phylogeny were removed (Table S1). This result also applied to other dietary classification schemes (Supplemental Figures S2 and S3).

For the anterior digastric muscle – the primary jaw opener in mammals - median normalized ACSA was slightly elevated in herbivorous omnivores than in carnivores, but with substantial overlap and data spread across all groups (Figure 2E). Herbivores show greater variance for the jaw opener, but there were no significant differences detected between diet groups (Table S1).

### Regression on dietary plant matter proportions

In addition to analyses considering diet as a categorical variable, we also treated percent dietary plant matter as a continuous variable. We used PGLS regressions to examine the relationship between muscle cross-sectional area and the percentage of plant matter in the diet.

As with the categorical analyses, the pterygoid complex had the strongest relationship between muscle proportion and percentage of plant matter in the diet, which showed a statistically significant, positive correlation – increased dietary plant matter was accompanied by proportionally larger pterygoid muscles (Figure 3A, Table S3). For the temporalis, we found a statistically significant negative relationship – species consuming more dietary plant matter had relatively smaller temporalis muscles (Figure 3B). We also found a statistically significant positive relationship between proportion of plant material and the relative size of the deep masseter complex (Figure 3C).

**Figure 3.**
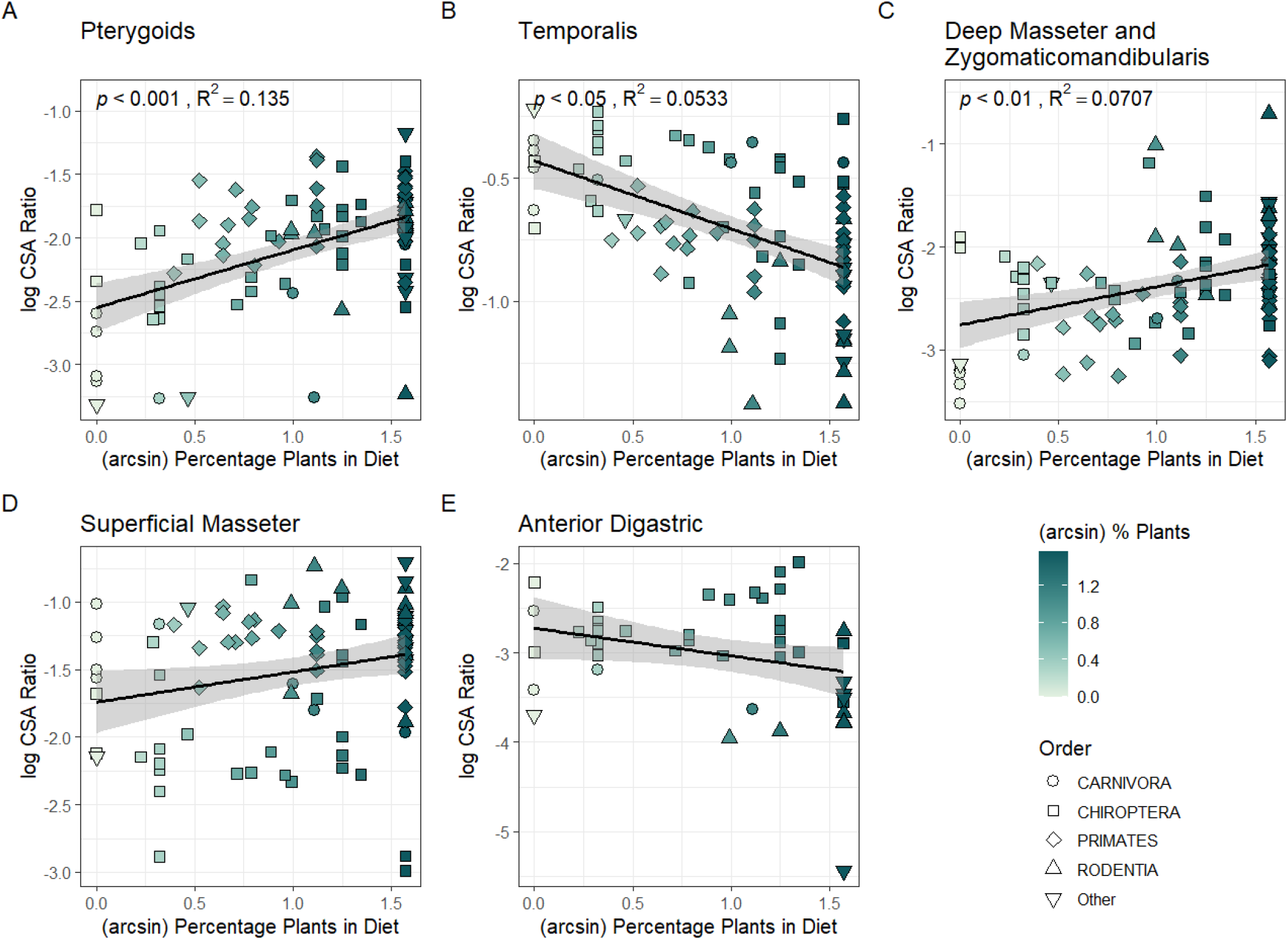
Scatter plots comparing the percentage of plant material in an animal’s diet to the proportional size of individual muscle phenotypes. Percentage plant matter data were linearized via an arcsine transformation prior to running phylogenetic regressions via PGLS and plotting in the figure above. While trend lines are shown for all muscles, PGLS regressions found a significant relationship between % plants and relative muscle size only for the pterygoid, temporalis and deep masseter muscles.

There was an apparent positive trend of the proportional size of superficial masseter increasing with the proportion of dietary plant matter. This trend was statistically significant in a standard regression analysis, (Table S4) but not in phylogenetic regression (Table S3). For anterior digastric, the regression line tended towards negative, however, phylogenetic regression did not recover a statistically significant relationship (Table S3).

### Phylogenetic principal components analysis

We used a phylogenetic PCA (pPCA) to visualize our muscle data in a multivariate context. Phylogenetic PC1 explained 56% of the variance in our dataset (Figure 4A) and was positively correlated with superficial masseter and pterygoid proportional size, and negatively correlated with temporalis and deep masseter complex proportional size (Figure 4B). Phylogenetic PC2 explained 28% of the dataset variance (Figure 4A), showing a positive correlation with temporalis proportional size and a negative correlation with the proportional size of all other jaw adductors (Figure 4B).

**Figure 4.**
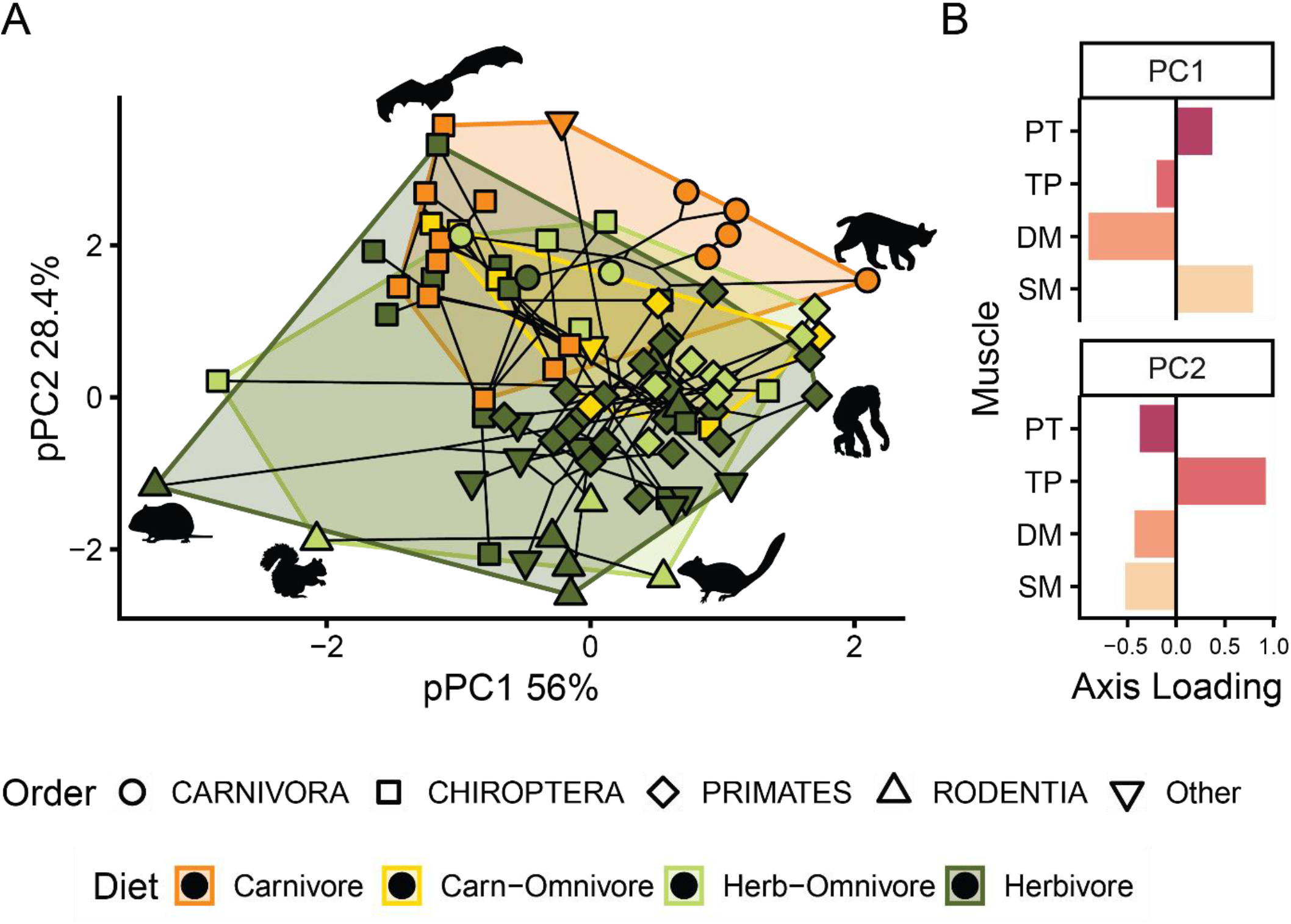
Phylogenetically corrected principal components analysis of muscle proportions (A), and loadings for each muscle across the first two principal component axes (B). The first and second principal component axes together explain 84.4% of the dataset variation with the first axis being strongly loaded by the relative size of the deep versus superficial masseter and pterygoid muscles (B) The secondary axis is mostly loaded by the relative size of the temporalis, compared to the other three muscle complexes (B). Herbivorous taxa tend to cluster at the positive end of PC1 and at the negative end of PC2, whereas carnivorous taxa tend to cluster towards the positive ends of PC1 and PC2. In A, points are color coded by diet, and shapes represent different taxonomic groupings. As shown by the convex hulls, there is considerable overlap between dietary groups. The phylogeny is overlaid onto the PCA plots (A, B), to illustrate the high degrees of homoplasy in muscle evolution and lack of phylogenetic structure in the data, corresponding to a strong dietary signal. Taxa at extremes of morphospace are illustrated with silhouettes and are (from the top, clockwise) *Noctilio albiventris, Lynx rufus, Pan troglodytes, Tamias striatus, Sciurus carolinensis*, and *Microtus arvalis*.

Carnivores tended to cluster at negative values of pPC1 and positive values of pPC2, whereas herbivores ranged towards the positive end of pPC1 and the negative end of pPC2 (Figure 4A). The two omnivorous groups occupied intermediate, overlapping regions. While there was some overlap among diets, there were clear trends along pPC1 and pPC2, consistent with dietary grouping.

## DISCUSSION

Despite a long-standing interest in the relationship between feeding behavior, diet, and mammalian jaw muscle morphology^3,4^, quantitative tests of these relationships have only been carried out relatively recently ^44–47,33,35,36^. Such tests have generally focused on specific mammalian groups, or dietary guilds^37^; and as a result, the identification of generalizable ecomorphological patterns across mammals has remained elusive. We gathered a taxonomically diverse dataset of jaw muscle cross-sectional area across major mammalian clades, showcasing multiple evolutionary transitions among carnivory, herbivory, and omnivory. We tested the relationships between diet and the relative proportion of the masseteric, temporalis, and pterygoid muscle complexes within a robust phylogenetic framework. Our results identify generalizable rules for mammalian jaw muscle evolution, primarily with the pterygoid complex (which predominantly consists of the medial pterygoid muscle) having the greatest explanatory power with respect to jaw function during mastication, and diet. Moreover, our data also reveal group-specific differences in how jaw muscle morphology relates to diet.

### Evolution of the pterygoids and grinding jaw movements in herbivores

Among the masticatory muscles, the pterygoids showed the strongest relationship between proportional muscle size (by proxy of ACSA), increasing in size with greater dependence on herbivory (proportion of plant matter consumed). This relationship persisted through phylogenetic corrections and was upheld irrespective of diet coding as a discrete or continuous variable (Figures 2A, 3A) and was also robust to variations in the number of dietary categories used (Supplemental Figures S2 and S3). The strong link between the pterygoids (specifically the medial pterygoid) and diet is an exciting result considering the extensive interest in pterygoid evolutionary development and the mechanical role of this complex in the evolution of transverse jaw movements in stem and crown mammals^48,49^. We hypothesize that transverse force application and work production of the medial pterygoid functionally underlies the association between elaboration of this muscle and herbivory across mammals. Herbivorous mammals require jaw translation to move the occlusal surfaces of their molars in a grinding manner to break down tough plant matter^20,29,50,51^. Grinding motions can be directed in the proal (anteroposterior) direction, as characteristic of rodents, and in the transverse (side-to-side) direction, as in most other herbivorous mammals (e.g., ungulates). Considering a single hemimandible, transverse translation may arise from several discrete motions, or combinations thereof, including pure medial translation, hemimandibular long-axis roll, and yaw rotation about a dorsoventral axis ^17–20^. Several muscles may drive these component motions, including deep masseter in primates ^52,53^, superficial masseter in rodents ^54–56^, and the medial belly of the pterygoid complex ^49^.

Medial pterygoid has a unique line of action among the jaw adductors, with fascicles running ventrally and laterally from the sphenoid skull base to insert onto the deep aspect of the hemimandible (Fig. 1B) ^4^. This line of action means that medial pterygoid can drive medial translation of the working hemimandible, yaw rotation and grinding occlusion during mastication. Medial pterygoid also plays a role in generating rostrally-directed movement ^3,57^, with hemimandibles either moving rostrally together for proal movement (most rodents) or moving individually to produce yaw (most non-rodent herbivores). During mastication, muscles are thought to either contribute to food processing (working-side) or stabilizing the mandible (balancing side)^58^. For the working-side, medial pterygoid forms part of the Triplet II muscle complex (with working-side superficial masseter and balancing-side temporalis) that contract concurrently to generate jaw yaw ^3,17,19,50,58,59^, by rotating the working side hemimandible laterally during the chewing power stroke. Therefore, medial pterygoid is clearly involved in most non-orthal jaw movements in mammals.

Biomechanical modeling, dental morphology, and dental occlusal patterns suggest that the first mammals to generate significant, medially-directed yaw during occlusion were stem therians ^6,19,20,60,61^. This group is characterized by a prominent, posteriorly-positioned angular process on the mandible, where medial pterygoid inserts. Angular process morphology covaries strongly with diet in mammals ^6^; the process is ventrally expanded in herbivores, displacing the ventral margin - and medial pterygoid insertion - further from the jaw joint, altering the line of action of the medial pterygoid and increasing torque for generating jaw yaw ^19^. Consequently, a relatively large medial pterygoid in extant herbivores (Fig. 2) likely represents a functional adaptation for increasing jaw yaw in order to effectively grind plant matter. Evolutionary reconstructions indicate that medial pterygoid was the most recent mammalian jaw muscle to evolve, possibly as recently as stem therians ^48,49,62^. Collectively, muscular and osteological evidence concur in suggesting a recent coevolution of medial pterygoid and angular process morphology as cornerstones in facilitating herbivory across mammals. The specialized nature of this anatomical and functional module for herbivory may also explain our observed patterns of dietary evolution.

The strong association between pterygoid complex size and diet provides a potential mechanistic explanation for observed patterns of dietary evolution across our sample. The ancestral dietary state in mammals appears to be carnivory^38^, but based on prior work on mammalian trophic evolution^41^ one might predict the majority of dietary transitions leading towards omnivory, *away* from both carnivory and herbivory^63^. However, our data reveal transitions *towards* more herbivorous diets to be much more frequent than the reverse (Supplemental Figure S1). The suite of adaptations that facilitate the evolution of herbivory – including the enlargement of the pterygoid musculature - form a series of bony, muscular and kinematic specializations which may constrain, or even channel dietary evolution. While previous studies found frequent transitions from herbivory to omnivory^41^, this pattern was largely driven by a high transition frequency in rodents, a clade that processes plant matter very differently from other herbivorous lineages meaning that they may experience a unique set of musculoskeletal constraints on dietary evolution^37^.

### Carnivores have large temporalis muscles for orthal, cutting bites

Our results for the temporalis substantiate an intuitively obvious yet long-lacking finding; that carnivorous mammals have temporalis muscles that are larger than herbivorous mammals (Figures 2 and 3). Temporalis acts to both elevate and retract the mandible, and a large temporalis muscle is generally associated with forceful vertical biting (mandible elevation), and cutting (mandible retraction)^4,64–67^. Whereas this result is consistent with prior empirical studies of the role of temporalis in vertical biting, they run counter to results from work on dietary associations based on bone proxies; Grossnickle (2020) found limited evidence of a relationship between diet and the height of the coronoid process, being the insertion site for the temporalis. This may be explained by temporalis being highly pennate with a tendinous insertion onto the coronoid process, but it may also reflect different functional signals in the soft vs hard tissue components (i.e., muscles vs. bones) of the feeding system, with muscles proving more informative indicators of diet ^32^.

That the temporalis signal was less clear-cut with respect to diet may be due to the precise functional role of this muscle. Forceful vertical biting, while probably most important for carnivorous taxa, is also functionally relevant to herbivores that carry or consume relatively large or hard food items (e.g., certain fruits or nuts) ^32,68–70^. Other herbivorous taxa have a relatively large temporalis, but have also altered the line of action of the muscle – more caudal, less dorsal - through changes in skull and jaw shape ^71^. In these taxa, the temporalis serves to horizontally retract the balancing side of the jaw during mastication, improving their efficiency for grinding plant material ^71^. The temporalis muscle may also be functionally relevant to gum-feeding marmosets that habitually gouge trees to stimulate the flow of exudate^72^. Outside of its role in feeding, relatively large temporalis muscles in males of sexually dimorphic taxa have been linked to generating high force for agonistic biting during aggressive male-male competition^73,74^.

### Dietary trends for the masseteric complex

The superficial masseter and deep masseteric complex, which is formed by the deep masseter and zygomaticomandibularis, are important jaw adductors in chewing. The deep masseteric complex is generally more orthally directed (dorsal skull origin, ventral mandibular insertion), whereas the superficial masseter often is directed more proally (rostral origin, caudal insertion) and, in concert with the medial pterygoid, responsible for proal jaw motion during occlusion^54,71,75^. As with the medial pterygoid, we also predicted masseteric proportional size to increase with herbivory. While our results showed such a trend, the signal did not withstand phylogenetic correction (Figures 2 and 3; Supplemental Figures 2 and 3).

There are several possible explanations for this weaker signal. First, variation in the proportional size of subunits in the masseteric complex is far greater than for temporalis and pterygoid across mammals, including among herbivores (Fig. 2B, D). Indeed, analyses of muscle mass proportions specifically in mammalian herbivores revealed considerable variation in the relative size of superficial masseter, deep masseter and zygomaticomandibularis; for example, whereas equids (zebra) had a prominent superficial masseter, this muscle was much smaller in ruminants (cows), where deep masseter was more prominent ^37^. Fascicle pennation angle was also more pronounced in the superficial masseter, whereas the deep masseter generally was more parallel-fibered^36^. Adaptations of the deep masseter to driving transverse jaw movements in primates ^76,77^ versus superficial masseter driving these same motions in rodents (see above) further exemplify the variable ecomorphology of the masseteric complex. The idea that masseteric muscle proportions map to diet in a many-to-one fashion ^78^ underscores the complexities associated with relating morphology of individual muscles to the emergent properties of the feeding system, like feeding mechanics and diet.

## Conclusion

Jaw muscle proportional size tracks dietary habits with strong to moderate affinity across mammals. In line with Maynard Smith & Savage’s (1959) hypothesis, we show that carnivores have enlarged temporalis muscles for generating orthal forces during biting whereas herbivores have enlarged medial pterygoid and superficial masseter muscles for generating rostral and transverse jaw translation, as well as jaw yaw during chewing and grinding. These patterns are broadly convergent across mammalian evolutionary radiations, highlighting the importance of soft-tissue adaptation of the masticatory apparatus in shaping the diversification of feeding strategies. The proportional size of the medial pterygoid, which was last to evolve in lineages leading into extant mammals, plays a key role in evolution away from carnivory, via omnivory, and into herbivory. This is consistent with the mechanical function of the medial pterygoid as well as superficial masseter in driving proal and transverse mandibular movements that are integral to molar grinding action for masticatory breakdown of tough plant matter in mammals.

## METHODS

### Specimen acquisition

Data on craniofacial muscle cross-sectional area were gathered for 91 mammal species with a diversity of diets (Figure 1A) (Data S1). Data for chiropterans and primates are from Santana (2018)^33^ and Taylor et al. (2025)^14^, respectively. Sampling of previously unpublished specimens focused on artiodactyls, perissodactyls, carnivorans, rodents, and lagomorphs. Specimens of these groups were salvaged as roadkill or acquired as donations from the Massachusetts Division of Fisheries and Wildlife (salvage permit 900C_20SAL, with updates). No animals were sacrificed for this study.

### Dietary Classifications

Dietary categorizations were based on the percentage of plant matter types requiring mastication (i.e., excluding soft fruits and nectar) that are consumed by a species ^6^ with supplemental evidence from other sources ^41–43^ (Data S1). We took two analytical approaches. First, we *discretized percent dietary plant matter* into a dietary scheme with four categories (carnivory, 0–15%; carnivorous omnivory, 15–50%; herbivorous omnivory, 50–85%; and herbivory, 85–100%). To test the sensitivity of our results to the chosen diet categorization scheme, we repeated analyses using two alternative schemes: a two-diet scheme (no omnivory) and a three-diet scheme (one omnivory group defined as having 15-85% plant diet) (see Supplemental Figure 2 and 3). Here carnivory and herbivory are broadly defined to include consumption of any animals (including insects) and plant material respectively. Second, we *treated percent dietary plant matter as a continuous variable*. We acknowledge that both approaches represent considerable simplifications of dietary schemes. However, they facilitate comparisons of species with high dietary specialization to those that show considerable seasonal fluctuations in diet, whilst also reducing the complexity of regression analyses and optimizing power for statistical analyses. Dietary evolution and transitions between diet types were modelled across our phylogeny using the *fitMK* function in the R package *phytools* (Supplemental Figure S1).

### Dissection and measurement approach

To measure the proportional sizes of each muscle phenotype in the feeding apparatus, we took morphological measurements from gross dissections of five muscles or their complexes: (1) the complex formed by lateral and medial pterygoids (hereafter, ‘pterygoids’), (2) the temporalis complex (which often consisted of discrete bellies), (3) the complex formed by deep masseter and zygomaticomandibularis, (4) the superficial masseter, and (5) the anterior digastric (a primary jaw depressor) (Figure 1B). For chiropteran and primate species, data collected and methods used were published previously ^14,33,79^. However, data were not available for all muscles across all specimens. Notably, due to the focus in primates on active force and passive stretch in the jaw-closing muscles, lateral pterygoid and anterior digastric data were unavailable for the primate sample. Moreover, the position of the pterygoid complex (Fig. 1B), deep to the mandible meant that dissection of this complex often was difficult and that the small lateral belly often could not be differentiated from the generally much larger medial belly. Therefore, we treated these two bellies as the pterygoid complex.

Jaw muscles were carefully dissected from both sides of each specimen, where available. Muscles were then blotted dry with tissue paper, weighed (muscle wet mass; M_M_) to the nearest 0.1 gram, sagittally bisected along their lines of action, and photographed with a metric ruler elevated to the cutting plane. In ImageJ (NIH) ^80^, we performed pixel-to-mm calibration of each photo using the ruler image and measured length (fascicle length: *L*_F_) and angular orientation (fascicle pennation; theta) with respect to collagenous insertions for at least three fascicles from each muscle belly as well as the muscle belly length (*L*_M_). Similar methods were used for the superficial masseter, medial pterygoid, and temporalis muscles in primates, with measurements taken *in situ* using digital calipers; acid digestion was used for the deep masseter ^30,31,36,81–83,73^. In chiropterans, myofiber lengths were taken following acid digestion, but because all species were small, their muscle fibers did not end intrafascicularly ^33^. Therefore, we are confident that the methodological difference did not affect the outcome of our analyses.

The degree to which muscles were appreciably pennate varied across taxonomical groups. Pennation was only consistently present in temporalis and superficial masseter. Consequently, and although pennation of the primate medial pterygoid often was substantial, we only factored pennation in calculations of cross-sectional area for those two muscles. For a sensitivity analysis on the effects of pennation angle, see the Supplemental Information (Supplemental Methods and Supplemental Figures S4 and S5). We calculated muscle physiological cross-sectional area (PCSA) as:

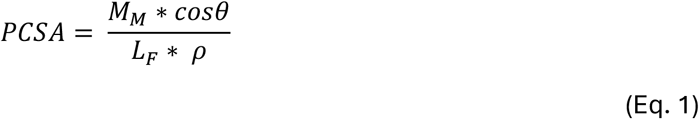

For all muscles we calculated anatomical cross-sectional area (ACSA) as:

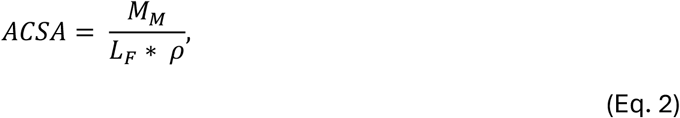

where cos*θ* is pennation angle, *L*_F_ is fascicle length, and *ρ* is the specific density of muscle ^84,85^.

To compare muscle proportions across species with different body sizes, we first normalized data on muscle cross-sectional area from each specimen by dividing PCSA or ACSA for each individual muscle by the total cross-sectional area of all jaw adductors (temporalis, superficial masseter, deep masseter and zygomaticomandibularis complex, and pterygoid complex). These proportional cross-sectional area values were then averaged across left and right sides for individuals, and across individuals of each species to generate grand-mean species averages. For downstream analyses, relative proportion data for each muscle phenotype were *ln* transformed, analogous to log-shape-ratios for linear morphometric measurements ^6,86^.

### Statistical testing

Following data inspection and outlier analysis using the *Rstatix* package, some taxa were excluded from further analyses either because (1) some muscles were incompletely sampled, such as in *Erethizon* (porcupine) where the pterygoid complex was not successfully dissected out, or (2) muscle phenotypes were absent, including a lack of temporalis in *Dipodomys* (kangaroo rat). While extreme muscle specializations, such as the one seen in *Dipodomys* represent real adaptations of the functional system, they would have potentially biased our dataset and obscured more subtle variation across our other taxa.

We analyzed the relationship between diet and muscle cross-sectional area using both discrete and continuous treatment of diet. For the four discrete diet categories, we used phylogenetic generalized least-squares (PGLS) analyses, followed by analysis of variance (ANOVA) to test for associations between measured muscle proportions and diet classification across species, while accounting for phylogenetic relatedness. Our phylogeny follows Upham et al (2019) ^87^, using a maximum clade credibility tree based on 500 trees from the posterior distribution and pruned to only include those taxa present in our dataset. We specified a correlation structure using Pagel’s lambda (λ) to model phylogenetic signal using the *nlme* package in R (v. 4.5.0). To assess pairwise differences among dietary categories, we conducted post hoc Tukey’s Honest Significant Difference (HSD) tests following each PGLS model using the *multcomp* package in R. For each trait’s PGLS model, we applied Tukey-adjusted multiple comparisons of means across all pairs of the four-level dietary classification. In all analyses, alpha was set to 0.05.

For our assessments of diet as a continuous variable, we used PGLS regressions to examine the relationship between muscle cross-sectional area and the percentage of plant matter in the diet. Trait values as species grand means were mapped and optimized to the tips of the phylogeny (Figure 1B) using the *comparative*.*data* function in the *caper* R package. These analyses incorporated the full variance-covariance matrix of the phylogeny. For each trait, a PGLS model was fitted using arcsine-transformed percent plant matter as the predictor, and model parameters were estimated under maximum likelihood for Pagel’s λ ^88^.

In addition to the univariate analyses presented above, we analyzed the relationship between proportional cross-sectional area of a given muscle phenotype with diet in a multivariate context. We tested for differences in jaw adductor proportional size across dietary groups, while accounting for phylogenetic structure, using a multivariate generalized least squares (GLS) model and multivariate analysis of variance (MANVOA) implemented in the R package mvMORPH ^89,90^. To visualize our data in a multivariate space, we conducted a phylogenetic principal components analysis (pPCA) on our set of jaw adductor muscle cross-sectional area measurements using the *phytools* package, implementing a Brownian motion model of trait evolution.

## Acknowledgements

Thanks to the coursework students at U. Mass. Lowell who assisted with PCSA data generation as part of a Curricular Undergraduate Research Experience sequence in the 2023 and 2024 editions of the course Form Feeds Function in Vertebrate Evolution. Data from this study formed part of a MS thesis for J. B.

## Data and code availability

All data required and code required to replicate the main findings of the paper are included as part of the supplemental information.

