## Supplemental Figures and Tables for "Dietary specializations are captured by jaw muscle proportions in mammals"

### Supplemental Information

#### Is pennation required, in addition to muscle mass and fascicle length, to accurately predict PCSA?

A secondary aim of our study was to determine how much variation in full PCSA is explained by the projection of fascicle length into muscle mass, and by muscle wet mass alone. Muscle proportional size is quantified using a variety of techniques, sparking recent confusion about the importance of muscle size in skeletal function. Techniques range from measuring muscle wet mass [1], to dividing muscle mass by the length of its fibers or fiber-bundles (fascicles) [2,3], to incorporating effects of both fiber/fascicle length and pennation [4–7]. The latter two approaches are collectively known as PCSA measurements and incorporate information about architecture (pennation) and sarcomere packing to better gauge muscle force production capacity. The architectural component of pennation is the degree of obliqueness of the attachment of fibers or fascicles in many muscles, relative to their line-of-action or force generation axis. Pennation has divergent effects on force by improving fiber packing and force generation from a geometric perspective [8], but also reduces force delivery along the line-of-action as a function of cosine to the pennation angle [9]. Debate about the mechanical importance of pennation has seen a recent upsurge [9,10], despite experimental evidence suggesting that a given muscle's peak tension can be estimated within 5% of the measured value based on muscle wet mass, average fiber/fascicle length, approximate pennation angle, and an estimate of specific tension estimated[11] (22.5 N/cm$^2$). We lack a broad comparative consensus on whether it is critical to incorporate pennation and/or fascicle length when measuring PCSA. In jaw systems, both superficial masseter and temporalis are suitable muscles for gauging the effect of measurement technique because they are large, have pronounced pennate architectures, are easily anatomically delineated, and relatively easy to dissect out [12]. Because our sample has a considerable range of animal body masses and muscle sizes, we additionally sought to determine if scaling influences how well muscle mass explains PCSA.

To compare cross-sectional areas calculated with and without pennation angle for the two consistently pennate muscles across our sample (superficial masseter and temporalis), we ran a GLS regression of CSA calculated as the projection of fascicle length into muscle mass (g/cm) against PCSA calculations incorporating pennation angle, muscle mass and fascicle length (cm$^2$) [11]. To determine how well muscle wet mass explains muscle PCSA, we recorded the regression slope, assuming that muscle cross-sectional area scales to muscle mass with an exponent of 0.67 [13,14]. The variance in full PCSA explained by the simpler measurement was greater than 99.0% for both the superficial masseter (Supplemental Figure S4A), and temporalis (Supplemental Figure S4B). Under isometry, the scaling exponent for muscle PCSA (cm$^2$) with respect to muscle wet mass (cm$^3$), should be 0.67 [13,14]. The regression slopes recovered from Ordinary Least-Squares analyses on log-transformed data were 0.74 for superficial masseter (Supplemental Figure S5A) and 0.70 for temporalis (Supplemental Figure S5B).

Although regressions showed almost all variation in PCSA computed using pennation angle was explained by PCSA computed without pennation angle, several taxa were outliers in this broad taxonomical comparison. These outliers may represent important physiological exceptions to an otherwise generalizable pattern and, since most outliers fall below the line of isometry

(Supplemental Figure S4A, B), their muscle architecture is likely specialized towards less pennation accompanied by greater mass or shorter length, with both of the latter traits representing excursions toward greater force production, possibly at the expense of contractile excursion. Since correlation does not equal causation, it is important to resist the temptation to generalize conclusions based on conjecture from sweeping analyses of large datasets. For instance, such generalizations have in the past led to the deeply rooted misconception that tendon stiffness is broadly conserved across mammals[15].

Muscle wet mass and PCSA scaled with positive allometry for both superficial masseter and temporalis where our measurements enabled a broad survey across disparate taxonomic mammal groups with considerable differences in body size (Supplemental Figure S5). There were both size- and taxonomic signals in the muscle mass/PCSA scaling relationships, with larger taxa, including ungulates, carnivores, and marsupials having proportionally heavier pennate muscles (superficial masseter and temporalis). This is consistent with the finding that fiber angle in pennate muscles increases with body size [5]. Conversely, smaller taxa, including bats and rodents tended to have greater PCSA for a given muscle mass with more acute pennation. The size and taxon dependency of these scaling relationships draw the practice of estimating muscle force production based on muscle wet mass into question (see also Martin et al 2020[16]), particularly when comparing animals of widely disparate sizes, and further enforce that both pennation, fascicle length, and possible fiber packing, matter[9] when attempting to gauge muscle force production capacity based on dissection measurements.

### Supplemental Figures

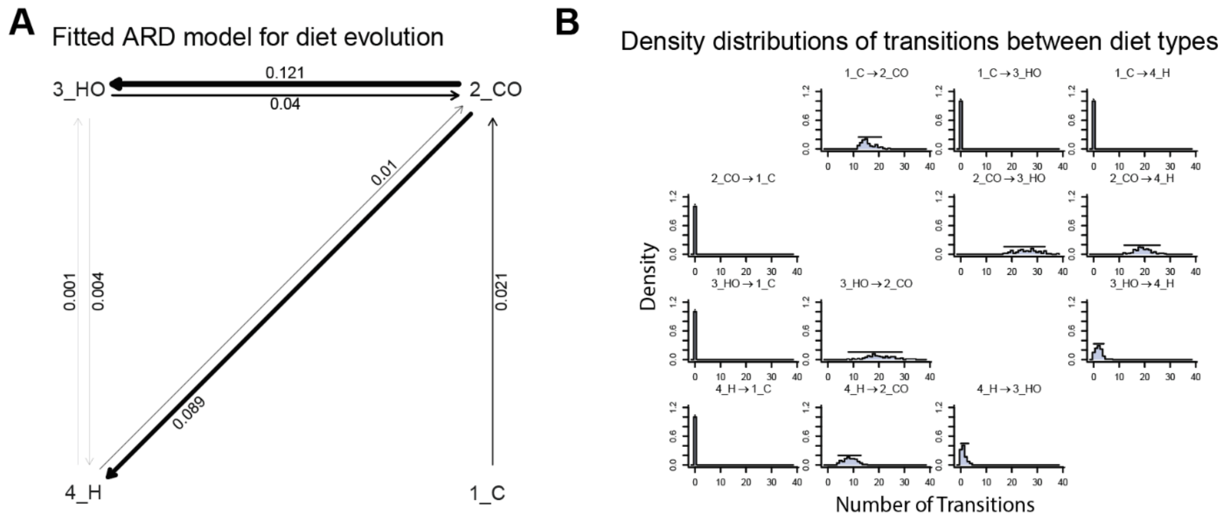


**Supplemental Figure S1**. Transitions between different dietary categories. (A) Fitted model of trait evolution, with estimated rates of transition from one diet to another - thicker arrows indicate more common transitions. (B) Density distributions of the estimated number of transitions between diet categories from a distribution of 100 stochastic character evolution maps. In both (A) and (B) the most frequent transitions are from carnivorous-omnivore to herbivorous-omnivore and herbivore. Overall, in our dataset, assuming the ancestral state for mammals was faunivorous, transitions towards more derived herbivorous diets are more common than the reverse.

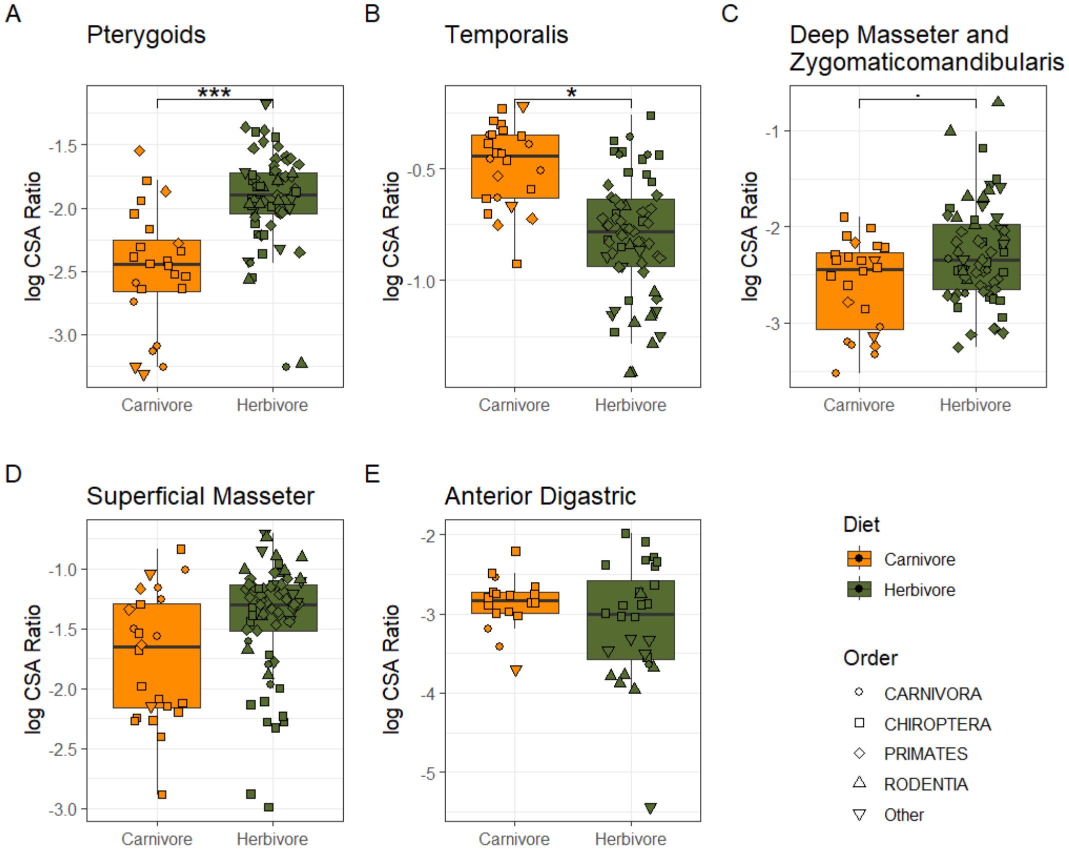


**Supplemental Figure S2**. Differences in muscle proportional size (separated in plot-space) by diet category, using a 2-diet scheme. Diets are carnivore (< 50% plant material), and herbivore (> 50% plant material). Muscles are the pterygoid (including medial and lateral pterygoid) (PT), temporalis (TP), zygomaticomandibularis and deep masseter (DM), superficial masseter (SM), and anterior digastric (AD). Significant differences between diet groups, analyzed in a PGLS framework, are shown for the temporalis and pterygoids, and a marginally significant result for the deep masseter.

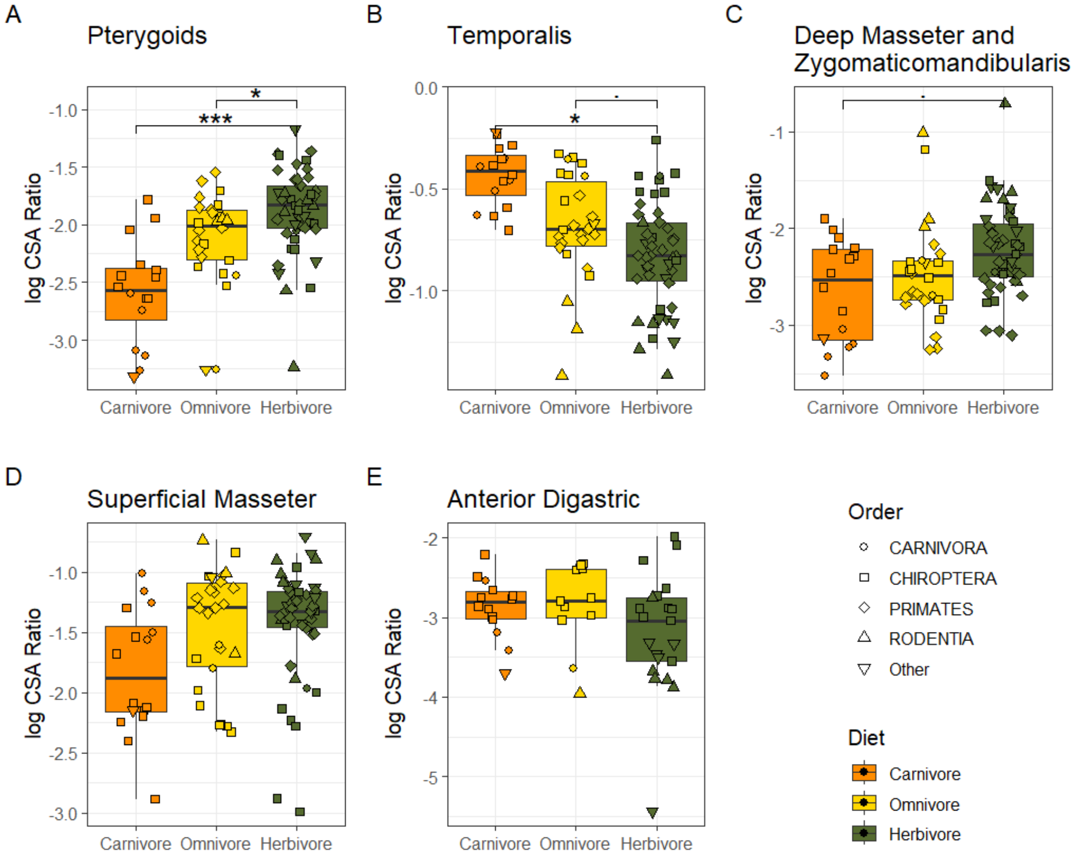


**Supplemental Figure S3**. Differences in muscle proportional size (separated in plot-space) by diet category using a 3-diet scheme. Diets are carnivore (< 15% plant material), omnivore (between 15% and 85% plant material) and herbivore (> 85% plant material). Muscles are the pterygoid (including medial and lateral pterygoid) (PT), temporalis (TP), zygomaticomandibularis and deep masseter (DM), superficial masseter (SM), and anterior digastric (AD). Significant differences between diet groups, analyzed in a PGLS framework, are shown for the temporalis and pterygoids, and a marginally significant result for the deep masseter.

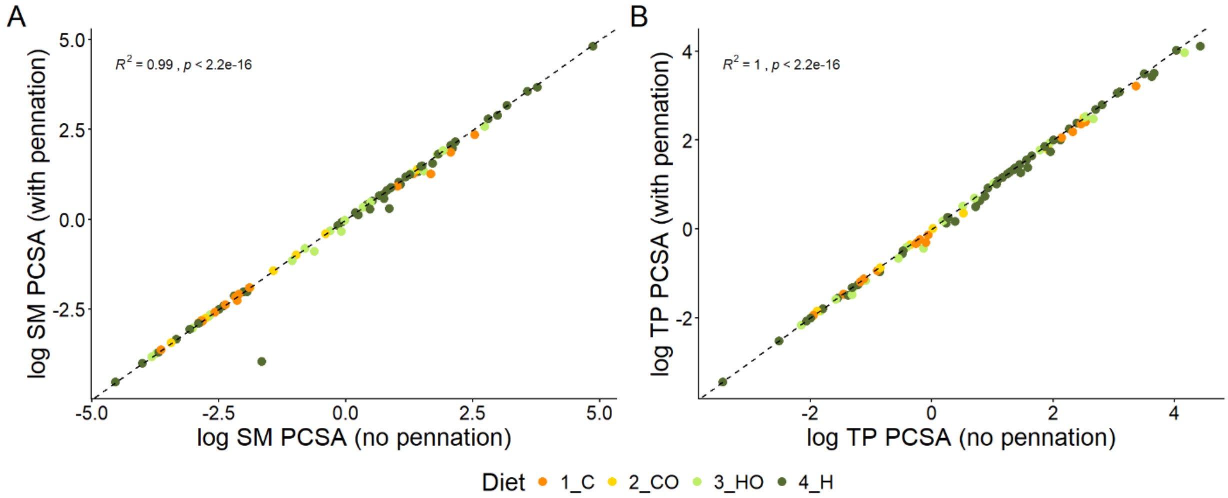


**Supplemental Figure S4**. Relationships between PCSA calculated using eq. 1. (y-axis, see text) and calculated as muscle mass divided by fascicle length (x-axis), without incorporation of fascicle pennation angle. This simpler measurement explains over 99% of the variation in the full PCSA dataset for both muscles.

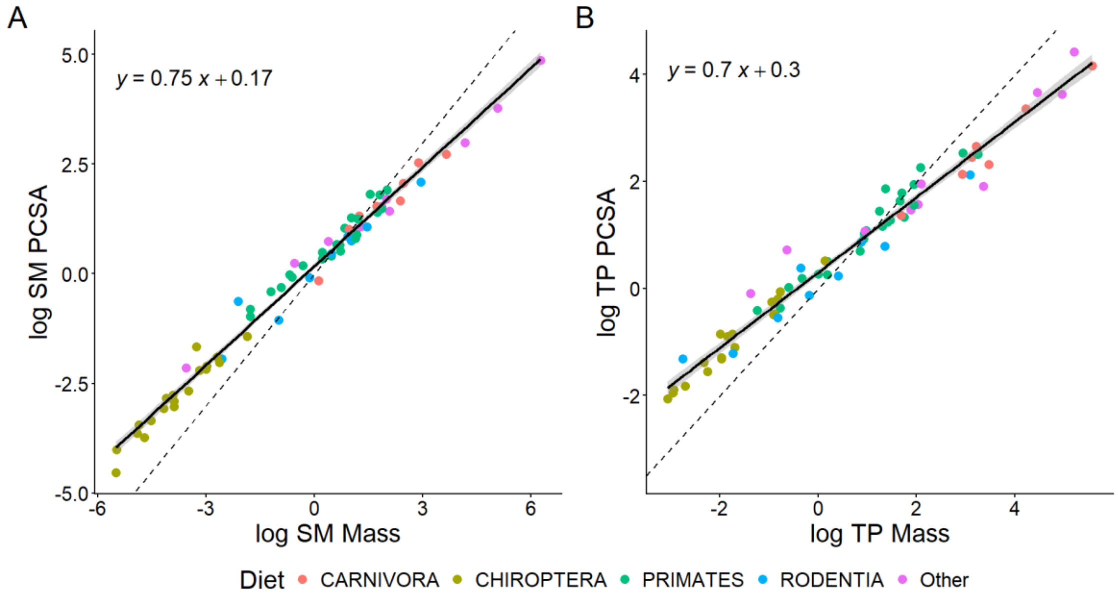


**Supplemental Figure S5**. Pennate muscle mass scales with positive allometry with respect to muscle PCSA (eq. 1) with a scaling exponent of 0.75 for superficial masseter (A) and 0.70 for temporalis (B).

### Supplemental Tables

Table S1. Post-hoc comparisons, following phylogenetic ANOVA

| Muscle | comparison | coefficients | sigma | tstat | pvalues |
|---|---|---|---|---|---|
| PT | 2_CO - 1_C | 0.137 | 0.137 | 0.994 | 0.75 |
| PT | 3_HO - 1_C | 0.285 | 0.131 | 2.18 | 0.127 |
| PT | 4_H - 1_C | 0.49 | 0.122 | 4.03 | 2.87e-4 |
| PT | 3_HO - 2_CO | 0.149 | 0.134 | 1.11 | 0.68 |
| PT | 4_H - 2_CO | 0.353 | 0.125 | 2.83 | 0.0236 |
| PT | 4_H - 3_HO | 0.205 | 0.103 | 1.98 | 0.191 |
| TP | 2_CO - 1_C | -0.0412 | 0.0804 | -0.512 | 0.956 |
| TP | 3_HO - 1_C | -0.0811 | 0.0792 | -1.02 | 0.733 |
| TP | 4_H - 1_C | -0.18 | 0.0739 | -2.44 | 0.069 |
| TP | 3_HO - 2_CO | -0.0399 | 0.0788 | -0.506 | 0.957 |
| TP | 4_H - 2_CO | -0.139 | 0.0737 | -1.89 | 0.231 |
| TP | 4_H - 3_HO | -0.0991 | 0.0623 | -1.59 | 0.381 |
| DM | 2_CO - 1_C | 0.0906 | 0.191 | 0.475 | 0.964 |
| DM | 3_HO - 1_C | 0.175 | 0.167 | 1.05 | 0.718 |
| DM | 4_H - 1_C | 0.333 | 0.152 | 2.19 | 0.123 |
| DM | 3_HO - 2_CO | 0.0843 | 0.184 | 0.458 | 0.967 |
| DM | 4_H - 2_CO | 0.243 | 0.169 | 1.44 | 0.471 |
| DM | 4_H - 3_HO | 0.158 | 0.132 | 1.2 | 0.62 |
| SM | 2_CO - 1_C | 0.082 | 0.181 | 0.452 | 0.969 |
| SM | 3_HO - 1_C | 0.0994 | 0.159 | 0.624 | 0.923 |
| SM | 4_H - 1_C | 0.123 | 0.145 | 0.845 | 0.83 |
| SM | 3_HO - 2_CO | 0.0174 | 0.175 | 0.0994 | 1 |
| SM | 4_H - 2_CO | 0.0407 | 0.161 | 0.253 | 0.994 |
| SM | 4_H - 3_HO | 0.0233 | 0.126 | 0.185 | 0.998 |
| AD | 2_CO - 1_C | -0.0944 | 0.204 | -0.463 | 0.966 |
| AD | 3_HO - 1_C | -0.0665 | 0.218 | -0.305 | 0.99 |
| AD | 4_H - 1_C | -0.203 | 0.19 | -1.07 | 0.704 |
| AD | 3_HO - 2_CO | 0.0279 | 0.264 | 0.106 | 1 |
| AD | 4_H - 2_CO | -0.109 | 0.237 | -0.459 | 0.967 |
| AD | 4_H - 3_HO | -0.137 | 0.197 | -0.693 | 0.897 |

Table S2. Post-hoc comparisons, following standard ANOVA

| Muscle | comparison | coefficients | sigma | tstat | pvalues |
|---|---|---|---|---|---|
| PT | 2_CO - 1_C | 0.288 | 0.172 | 1.68 | 0.333 |
| PT | 3_HO - 1_C | 0.532 | 0.136 | 3.9 | 9.92E-04 |
| PT | 4_H - 1_C | 0.707 | 0.115 | 6.16 | 9.78E-08 |
| PT | 3_HO - 2_CO | 0.244 | 0.169 | 1.45 | 0.463 |
| PT | 4_H - 2_CO | 0.419 | 0.152 | 2.76 | 0.0334 |
| PT | 4_H - 3_HO | 0.174 | 0.11 | 1.59 | 0.382 |
| TP | 2_CO - 1_C | -0.154 | 0.106 | -1.46 | 0.455 |
| TP | 3_HO - 1_C | -0.302 | 0.0837 | -3.61 | 0.00258 |
| TP | 4_H - 1_C | -0.393 | 0.0705 | -5.58 | 2.21E-06 |
| TP | 3_HO - 2_CO | -0.148 | 0.104 | -1.43 | 0.475 |
| TP | 4_H - 2_CO | -0.239 | 0.0932 | -2.57 | 0.0547 |
| TP | 4_H - 3_HO | -0.0911 | 0.0676 | -1.35 | 0.526 |
| DM | 2_CO - 1_C | 0.129 | 0.214 | 0.604 | 0.928 |
| DM | 3_HO - 1_C | 0.208 | 0.17 | 1.22 | 0.604 |
| DM | 4_H - 1_C | 0.427 | 0.143 | 2.98 | 0.0184 |
| DM | 3_HO - 2_CO | 0.0787 | 0.21 | 0.374 | 0.981 |
| DM | 4_H - 2_CO | 0.297 | 0.189 | 1.57 | 0.39 |
| DM | 4_H - 3_HO | 0.219 | 0.137 | 1.6 | 0.377 |
| SM | 2_CO - 1_C | 0.261 | 0.211 | 1.24 | 0.597 |
| SM | 3_HO - 1_C | 0.396 | 0.168 | 2.36 | 0.0891 |
| SM | 4_H - 1_C | 0.409 | 0.141 | 2.9 | 0.0233 |
| SM | 3_HO - 2_CO | 0.135 | 0.207 | 0.65 | 0.912 |
| SM | 4_H - 2_CO | 0.148 | 0.187 | 0.795 | 0.852 |
| SM | 4_H - 3_HO | 0.0137 | 0.135 | 0.101 | 1 |
| AD | 2_CO - 1_C | 0.0254 | 0.351 | 0.0726 | 1 |
| AD | 3_HO - 1_C | 0.00293 | 0.286 | 0.0103 | 1 |
| AD | 4_H - 1_C | -0.311 | 0.213 | -1.46 | 0.461 |
| AD | 3_HO - 2_CO | -0.0225 | 0.388 | -0.0581 | 1 |
| AD | 4_H - 2_CO | -0.336 | 0.337 | -0.997 | 0.744 |
| AD | 4_H - 3_HO | -0.314 | 0.27 | -1.16 | 0.644 |

Table S3. Phylogenetic regression results

| Muscle | Fv | pv | R2 | La |
|---|---|---|---|---|
| PT | 14.74 | 2.339e-4 | 0.135 | 0.8607 |
| TP | 5.955 | 0.0167 | 0.05331 | 0.892 |
| DM | 7.691 | 0.006788 | 0.07067 | 0.5802 |
| SM | 0.15 | 0.6995 | -0.00975 | 0.6392 |
| AD | 1.77 | 0.1903 | 0.01682 | 0.926 |

Table S4. Standard regression results

| Muscle | pv | R2 |
|---|---|---|
| PT | 2.99E-07 | 0.2618 |
| TP | 2.22E-07 | 0.2667 |
| DM | 1.97E-04 | 0.148 |
| SM | 0.02526 | 0.05623 |
| AD | 0.05866 | 0.07888 |
